# ADMET-AI: A machine learning ADMET platform for evaluation of large-scale chemical libraries

**DOI:** 10.1101/2023.12.28.573531

**Authors:** Kyle Swanson, Parker Walther, Jeremy Leitz, Souhrid Mukherjee, Joseph C. Wu, Rabindra V. Shivnaraine, James Zou

## Abstract

**Summary:** The emergence of large chemical repositories and combinatorial chemical spaces, coupled with high-throughput docking and generative AI, have greatly expanded the chemical diversity of small molecules for drug discovery. Selecting compounds for experimental validation requires filtering these molecules based on favourable druglike properties, such as Absorption, Distribution, Metabolism, Excretion, and Toxicity (ADMET). We developed ADMET-AI, a machine learning platform that provides fast and accurate ADMET predictions both as a website and as a Python package. ADMET-AI has the highest average rank on the TDC ADMET Benchmark Group leaderboard, and it is currently the fastest web-based ADMET predictor, with a 45% reduction in time compared to the next fastest ADMET web server. ADMET-AI can also be run locally with predictions for one million molecules taking just 3.1 hours.

**Availability and Implementation:** The ADMET-AI platform is freely available both as a web server at admet.ai.greenstonebio.com and as an open-source Python package for local batch prediction at github.com/swansonk14/admet_ai (also archived on Zenodo at doi.org/10.5281/zenodo.10372930). All data and models are archived on Zenodo at doi.org/10.5281/zenodo.10372418.

## Introduction

The curation of large chemical repositories and advances in algorithms and computational hardware have significantly expanded the scale of *in silico* drug discovery campaigns. Structure-based virtual screening, pharmacophore modelling and, in particular, high-throughput molecular docking (HTMD) have been used to screen over a billion molecules against therapeutic targets of interest^1^. More recently, generative artificial intelligence (AI) approaches have been developed to design a vast array of compounds that are optimised for a particular therapeutic effect^2^. Both HTMD and generative AI approaches yield a large number of potential hits with high efficacy, but many of these hits are not ideal for therapeutic development because they do not possess druglike properties^3^. As a result, rapid and accurate screening of these hits for ideal druglike properties is critical for advancing molecules with higher probabilities of success^4^. Specifically, for a small molecule to be at the centre of a therapeutic strategy and proceed from discovery to clinical trials, the compound must possess optimal Absorption, Distribution, Metabolism, Excretion, and Toxicity (ADMET) properties.

Here, we have developed ADMET-AI, a simple, fast, and accurate platform for ADMET property prediction that includes both a web interface and a Python package for local prediction (**Figure 1**). ADMET-AI uses a graph neural network called Chemprop-RDKit (**Figure 1A**), which was trained on 41 ADMET datasets from the Therapeutics Data Commons (TDC). ADMET-AI surpasses existing ADMET prediction tools in terms of speed and accuracy (**Figure 1B**,**C**). Moreover, it provides additional useful features such as local batch prediction (**Figure 1D**) and contextualised ADMET predictions using a reference set of approved drugs; these features are missing in most of the current ADMET prediction tools. We make ADMET-AI freely available (**Figure 1E**) and open-source as a resource to aid in the evaluation of large-scale chemical libraries for druglike compounds.

**Figure 1:**
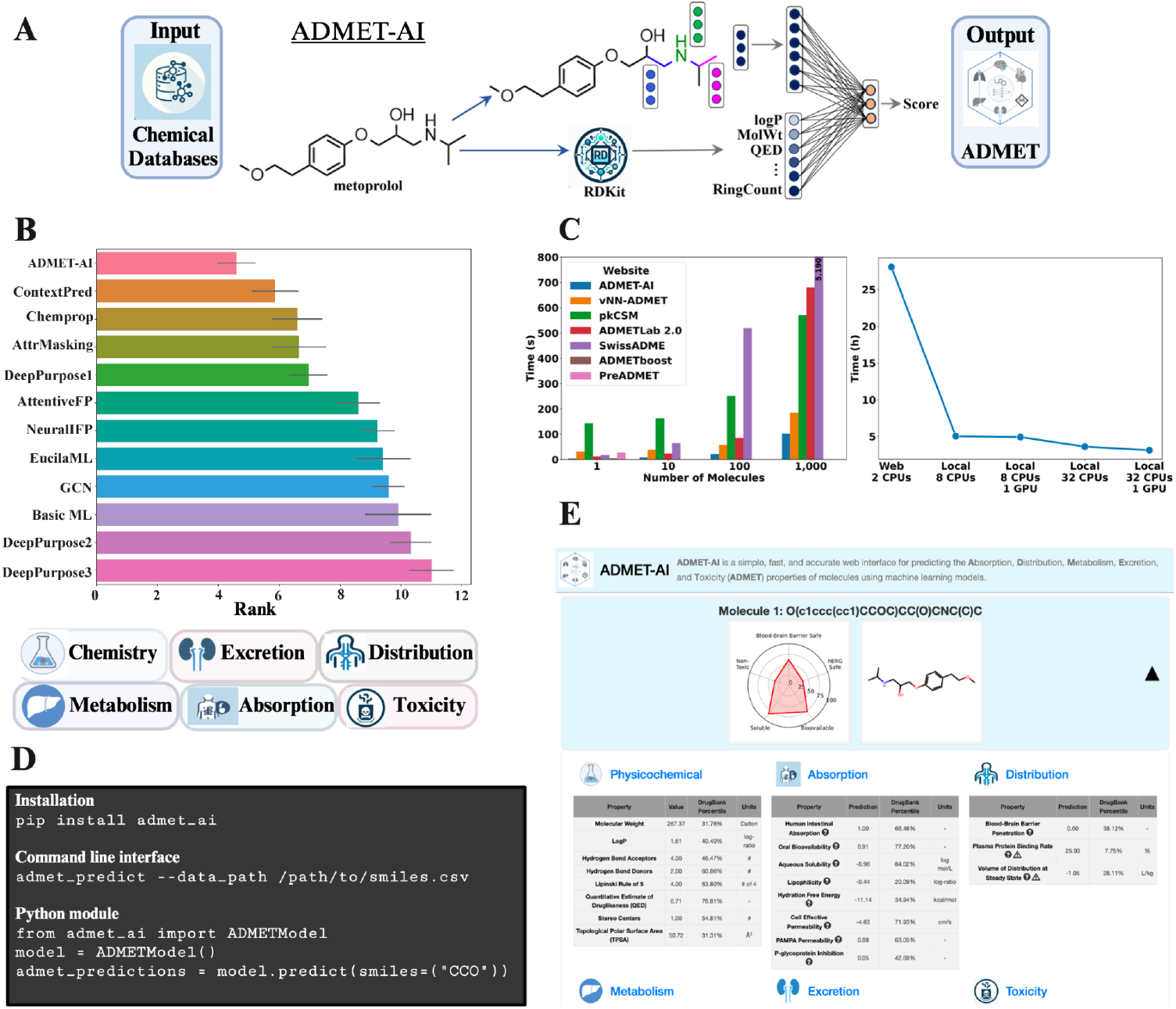
Overview of ADMET-AI. **A)** An illustration of training an ADMET-AI graph neural network Chemprop-RDKit model. **B)** The overall rank of ADMET-AI models on the Therapeutics Data Commons ADMET leaderboard of 22 ADMET datasets. Representative overall categories predicted by ADMET-AI are shown below. **C**) The computational efficiency of ADMET-AI. *Left panel*, the time (in seconds, median of three trials) for the ADMET-AI web server and other common ADMET web servers to make predictions on 1, 10, 100, or 1,000 molecules from the DrugBank reference set (median of three trials). ADMETboost and PreADMET are limited to one molecule. Since pkCSM and SwissADME are limited to 100 and 200 molecules, respectively, their 1,000 molecule times are computed as 100 molecule time x 10. *Right panel*, the time (in hours, median of three trials) for the ADMET-AI web server and various hardware configurations running the ADMET-AI local command line tool to make predictions on 1 million molecules from the DrugBank reference set (1,000 copies of the 1,000 molecule DrugBank set). Since the ADMET-AI web server is currently limited to 1,000 molecules, its 1 million molecule time is computed as 1,000 molecule time x 1,000. **D)** Commands needed to install and run the local version of ADMET-AI, either as a command line tool or as a Python module. **E)** Predictions displayed on the ADMET-AI website (admet.ai.greenstonebio.com).

## Model development

### Data

ADMET-AI uses 41 ADMET datasets (10 regression, 31 classification) from the Therapeutics Data Commons^5^ (TDC, v0.4.1). Of those 41 datasets, 22 datasets (9 regression, 13 classification) form the TDC ADMET Benchmark Group leaderboard, which enables a side-by-side comparison of ADMET-AI compared to other ADMET prediction models. Additionally, from the 41 ADMET datasets, two multi-task datasets were created, one containing all 10 regression datasets and one containing all 31 classification datasets. This enables the training of just two models (one regression, one classification) that cover all 41 ADMET properties. Summary statistics for the ADMET datasets are listed in **Supplementary Table 1**, and further details on the preparation of the datasets are in the supplementary methods.

### Model

ADMET-AI uses a deep learning model architecture for ADMET prediction called Chemprop-RDKit^6^ (available in Chemprop, v1.6.1). Chemprop-RDKit consists of a graph neural network called Chemprop that is augmented with 200 physicochemical molecular features that are computed by the cheminformatics package RDKit^7^ (v2023.3.3). The Chemprop graph neural network computes simple features for each atom (*e*.*g*., atom type) and each bond (*e*.*g*., bond type) and then runs several steps of message passing with neural network layers to aggregate atom and bond information across the molecule to build a single representation of the whole molecule. This representation is then concatenated with the 200 RDKit features, and the combined representation is passed through several feed-forward neural network layers to predict one or more endpoints, depending on the dataset.

For each TDC dataset, one Chemprop-RDKit model was trained for each of the five train/validation/test splits, and the average performance across the five test splits for that dataset was recorded. At inference time, for each dataset, the average prediction is computed from an ensemble of models containing the five models trained on the five splits of that dataset. Ensembling is used since it can overcome incorrect biases in individual models and thereby improve the performance and robustness of the ensemble^8^.

### Results

On the 22 datasets in the TDC ADMET Benchmark Group leaderboard, ADMET-AI has the best average rank among all models that were evaluated on these datasets (**Figure 1B, Supplementary Figure 1**). While other models are accurate on only a few ADMET properties, ADMET-AI is the best on average across all ADMET properties, thereby making it ideal for comprehensive ADMET analysis.

Across all 41 TDC ADMET datasets, ADMET-AI models trained on each dataset individually (“single-task”) had overall strong performance, with an R^2^ greater than 0.6 for five of the ten regression datasets and an AUROC greater than 0.85 for 20 of the 31 classification datasets (**Supplementary Figure 2**). ADMET-AI models trained on the two multi-task datasets achieved very similar performance (**Supplementary Figure 2**). Notably, the two multi-task models (one for regression, one for classification) make predictions significantly faster than the 41 single-task models. Therefore, the multi-task models were deployed in the ADMET-AI web server.

Complete performance results are in **Supplementary Table 1**.

### ADMET-AI Web Server

The ADMET-AI web server was built using Flask^9^ (v2.3.2), which is a Python-based micro web framework. ADMET-AI possesses an intuitive user interface to display the predicted ADMET properties. Additionally, each predicted ADMET value is contextualised via a comparison to a reference set of molecules from the DrugBank as described below. The ADMET-AI models can also be run locally for high-throughput ADMET prediction. Both as a web server and as a local package, ADMET-AI is significantly faster than alternate ADMET prediction tools.

### DrugBank Reference

ADMET predictions can be difficult to interpret in isolation and can be better evaluated within the context of drugs with a similar therapeutic indication. As such, ADMET-AI incorporates a point of reference by comparing each query molecule to a set of approved molecules. This unique feature provides useful context for interpreting ADMET predictions and guides the user to the range of ADMET properties needed for their drug discovery project. ADMET-AI, therefore, curated a list of 2,579 drugs from the DrugBank^10^ (v5.1.10) that have obtained regulatory approval as a reference set (**Supplementary Table 2**). The ADMET-AI multi-task models were applied to make predictions on these molecules for all ADMET endpoints. For any molecule queried on the ADMET-AI website, ADMET-AI computes the percentile of that molecule’s ADMET predictions relative to this DrugBank reference. Furthermore, since it is known that different classes of drugs have different ADMET requirements (*e*.*g*., acceptable toxicity for antineoplastics is much higher than that for antibiotics), the ADMET-AI website, therefore, allows the user to select an appropriate subset of the DrugBank reference based on Anatomical Therapeutic Chemical (ATC) codes^11^ (common ATC codes are shown in **Supplementary Figure 3**) to use as the reference set for computing percentiles to add perspective during evaluation.

### User Experience

Users access the web interface of ADMET-AI at admet.ai.greenstonebio.com as shown in **Figure 1E**. There are three options to input molecules: (1) by entering SMILES in a text box with each SMILES on a line, (2) by uploading a CSV file containing multiple SMILES, or (3) by drawing the structure of the molecule using an interactive tool that converts the drawing to a SMILES. Users can make predictions on up to 1,000 molecules at a time. Next to the SMILES input, the user can create the DrugBank reference set by selecting an ATC code or by using the default of all DrugBank-approved molecules.

Completed ADMET predictions, along with physicochemical properties computed by RDKit, are displayed on the website for up to 25 molecules (**Figure 1E**), with predictions for the remaining molecules available for download. For each predicted molecule, a radial plot is shown as a quick summary of key components of druglikeness, including hERG toxicity (*i*.*e*., potential for cardiotoxicity and arrhythmias), blood-brain barrier penetration (*i*.*e*., potential for CNS effects), solubility, oral bioavailability (*i*.*e*., ease of drug delivery), and potential for toxicity. The complete set of ADMET predictions for the molecule is shown in a set of tables, and each prediction is paired with the percentile of that prediction with respect to the DrugBank-approved reference set. For regression properties, the predicted values are displayed in the same units as in the dataset used to train the model (*e*.*g*., half life (t_1/2_) is predicted in terms of hours). For binary classification properties, the predicted values are the probability that the molecule has the property (*e*.*g*., the probability of blood-brain barrier (BBB) penetration). For a simultaneous comparison of multiple molecules, ADMET-AI displays a summary scatter plot with two user-selected ADMET properties (one on each axis) as compared to the DrugBank reference set.

### Local ADMET Prediction

The ADMET-AI website enables fast and accurate ADMET prediction with no installation or computational experience required. ADMET-AI is also available as a Python package (github.com/swansonk14/admet_ai) for local ADMET prediction using the same multi-task ADMET models that power the ADMET-AI website. The single-task models are also available for download and use. The local version of ADMET-AI includes a command line tool called `admet_predict` that allows users to make predictions on massive datasets, such as those originating from large docking campaigns or generative AI models, that would exceed the computational capacity of the web server (**Figure 1D**). The ADMET models can also be incorporated directly within a user’s local computational pipeline by importing the ADMET-AI models in just a few lines of Python code (**Figure 1D**). The local version of ADMET-AI has no limit on the number of molecules that can be run through the model, and it automatically uses a GPU, if available, and multiple CPUs to increase prediction speed. The local version is also ideal for users who are privacy conscious and want to avoid putting their proprietary compounds through a web server, although it should be noted that the ADMET-AI web server does not store information on any molecules that are queried.

### Comparison to Alternatives: Speed

While there are several ADMET prediction web servers currently available, ADMET-AI provides a powerful combination of accuracy, speed, and flexibility along with unique features such as the DrugBank reference set and local prediction. Additionally, ADMET-AI is fully open source, making it easy to build upon and extend.

One of the most important qualities of an ADMET tool is the speed with which it can predict the ADMET properties of molecules. This is increasingly important as drug discovery projects rapidly grow in scale. To that end, the speed of ADMET-AI was compared to six other web servers that predict a wide range of ADMET properties: SwissADME^12^, ADMETLab 2.0^13^, pkCSM^14^, vNN-ADMET^15^, ADMETboost^16^, and PreADMET^17,18^. Each server was used to make ADMET predictions for 1, 10, 100, or 1,000 molecules from the DrugBank reference (**Supplementary Table 3**), and the median of three trials was recorded. ADMET-AI is the fastest web server for any number of molecules, with a 45% reduction in time over the next best server for 1,000 molecules (**Figure 1C**, *left panel*).

For truly large-scale analysis, local prediction is preferable to take advantage of greater CPU parallelism and GPU availability. To illustrate this, local versions of ADMET-AI with 8 or 32 CPU cores and with or without a GPU were used to make predictions on one million molecules from the DrugBank reference (1,000 copies of the 1,000 molecule DrugBank set). The local prediction speed was compared to the speed of the ADMET-AI web server, which uses two CPU cores and no GPU. Local ADMET-AI prediction with 32 CPU cores and a GPU reduces the ADMET prediction time from 28.1 hours with the web server to 3.1 hours locally, representing a further 89% reduction in time (**Figure 1C**, *right panel*). Even a simple setup of 8 CPU cores and no GPU can make one million predictions in just five hours. Therefore, ADMET-AI enables fast, large-scale ADMET prediction.

## Conclusion

ADMET-AI is a simple, fast, and accurate platform for ADMET prediction. ADMET-AI uses a Chemprop-RDKit graph neural network for ADMET prediction that is currently the most accurate model on average across the Therapeutics Data Commons ADMET leaderboard. The ADMET-AI website is the fastest web-based ADMET predictor with a 45% reduction in time compared to the next fastest ADMET web server. The ADMET-AI website uniquely provides context for ADMET predictions by comparing predictions on input molecules to predictions on approved drugs from the DrugBank, which can optionally be filtered to a specific drug category by ATC code. In addition to the web interface, the ADMET-AI platform includes a Python package that can be applied locally either as a command line tool for large-scale evaluation or as a Python module for use within other drug discovery tools (e.g., generative AI models). ADMET-AI is an open-source, free, and easy-to-use platform that can serve as a powerful drug discovery tool for identifying compounds with favourable ADMET profiles for further development.

## Author Contributions

Conceptualization [K.S., P.W., J.L., S.M., J.C.W., R.V.S., J.Z.], methodology [K.S., P.W., R.V.S, J.Z.], software [K.S., P.W.], supervision [J.C.W., R.V.S., J.Z.], visualisation [K.S., P.W., R.V.S.], writing – original draft [K.S., R.V.S., J.Z.].

## Acknowledgments

We thank Jonathan Stokes and Gary Liu for providing design ideas and feedback for the ADMET-AI website. We thank Kirk Swanson for testing and providing feedback for the ADMET-AI website. We thank Kexin Huang for help posting the ADMET-AI results to the Therapeutics Data Commons leaderboard. This work was supported by the Knight-Hennessy Scholarship [to K.S.] and by the National Institutes of Health [R01 HL163680 and R01 HL171102 to J.C.W.].

## Supplementary Methods

### TDC Datasets

ADMET-AI models were trained on three subsets of the datasets from the Therapeutics Data Commons (TDC). Details of each subset are described below.

#### TDC Single-Task

The TDC Single-Task subset contains 41 ADMET datasets. It includes all ADMET datasets in the TDC except “hERG_Karim” and “herg_central,” which are redundant with the “herg” dataset, and “ToxCast,” which has 617 endpoints and is therefore challenging to interpret and to display on a web server. Among the 30 datasets, 10 are regression datasets and 31 are binary classification datasets. Each dataset was split into train, validation, and test sets using an 80%, 10%, 10% random split with five different random seeds. This makes it possible to train an ensemble of five models, each of which has seen a wide variety of molecular scaffolds, thereby improving generalisation of the models compared to using a scaffold split but at the potential cost of overly optimistic test set performance due to scaffold sharing between the train and test sets. For the regression datasets, the metrics were MAE and R^2^ (coefficient of determination), and for the classification datasets, the metrics were AUROC and AUPRC.

#### TDC Leaderboard

The TDC Leaderboard subset contains the 22 ADMET datasets in the TDC ADMET Benchmark Group (a subset of the 41 datasets in TDC Single-Task). Nine of the datasets are regression datasets and 13 of the datasets are binary classification datasets. The TDC maintains a leaderboard for the TDC Leaderboard datasets that ranks models according to their performance on a held-out test set for each dataset. This makes it possible to directly compare ADMET-AI’s model against other state-of-the-art ADMET prediction tools. As per the TDC leaderboard guidelines, five models were trained, each one using a Murcko scaffold split with roughly 87.5% training data and 12.5% validation data created with different random seeds, and all models were evaluated using a single scaffold-split test set provided by the TDC. Scaffold splits represent the challenging but realistic scenario where the model is trained on molecules with certain chemical scaffolds but must generalise to new molecules with different scaffolds (e.g., new classes of drugs). For each dataset, the metric provided by the TDC is either mean absolute error (MAE) or Spearman’s rank correlation coefficient (Spearman) for regression datasets and is either area under the receiver operating characteristic curve (AUROC) or area under the precision-recall curve (AUPRC) for binary classification datasets. The leaderboard results reported here are as of October 4, 2023. Only models that are evaluated on all 22 datasets in the leaderboard are included.

#### TDC Multi-Task

The TDC Multi-Task subset contains two multi-task datasets built from TDC Single-Task. One dataset contains all 10 regression datasets (22,100 unique compounds), and the other contains all 31 binary classification datasets (35,774 unique compounds). The datasets were merged by matching SMILES so that any molecule that appears in multiple TDC Single-Task datasets appears as a single row in the multi-task dataset containing its SMILES and its endpoint values from every dataset it appears in. For datasets that the molecule does not appear in, the molecule’s endpoint is filled in as a null value, and that endpoint is ignored during training and has no effect on the model’s learning. The same random splitting scheme and metrics as for the TDC Single-Task datasets were used.

**Supplementary Figure 1:**
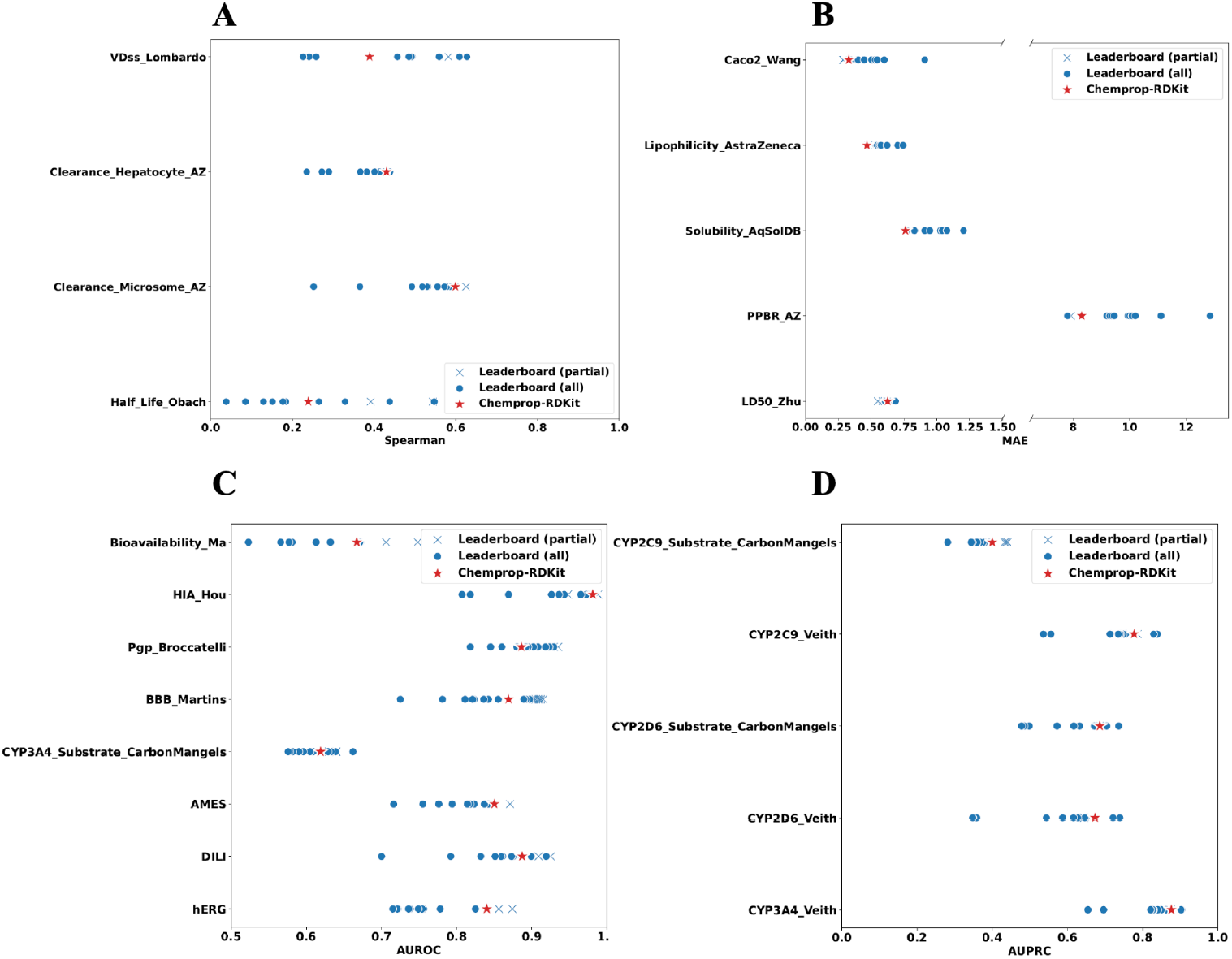
TDC Leaderboard Results. Performance of ADMET-AI (red star) compared to other models (blue dots and crosses) on the TDC Leaderboard datasets according to four different metrics. Blue dots indicate models evaluated on all 22 TDC Leaderboard datasets while blue crosses indicate models evaluated on only some of the datasets (often just one). **A–B)** Performance on the regression datasets using either **(A)** Spearman rank correlation coefficient or **(B)** mean absolute error (MAE). **C–D)** Performance on the classification datasets using either **(C)** area under the receiver operating characteristic curve (AUROC) or **(D)** area under the precision-recall curve (AUPRC).

**Supplementary Figure 2:**
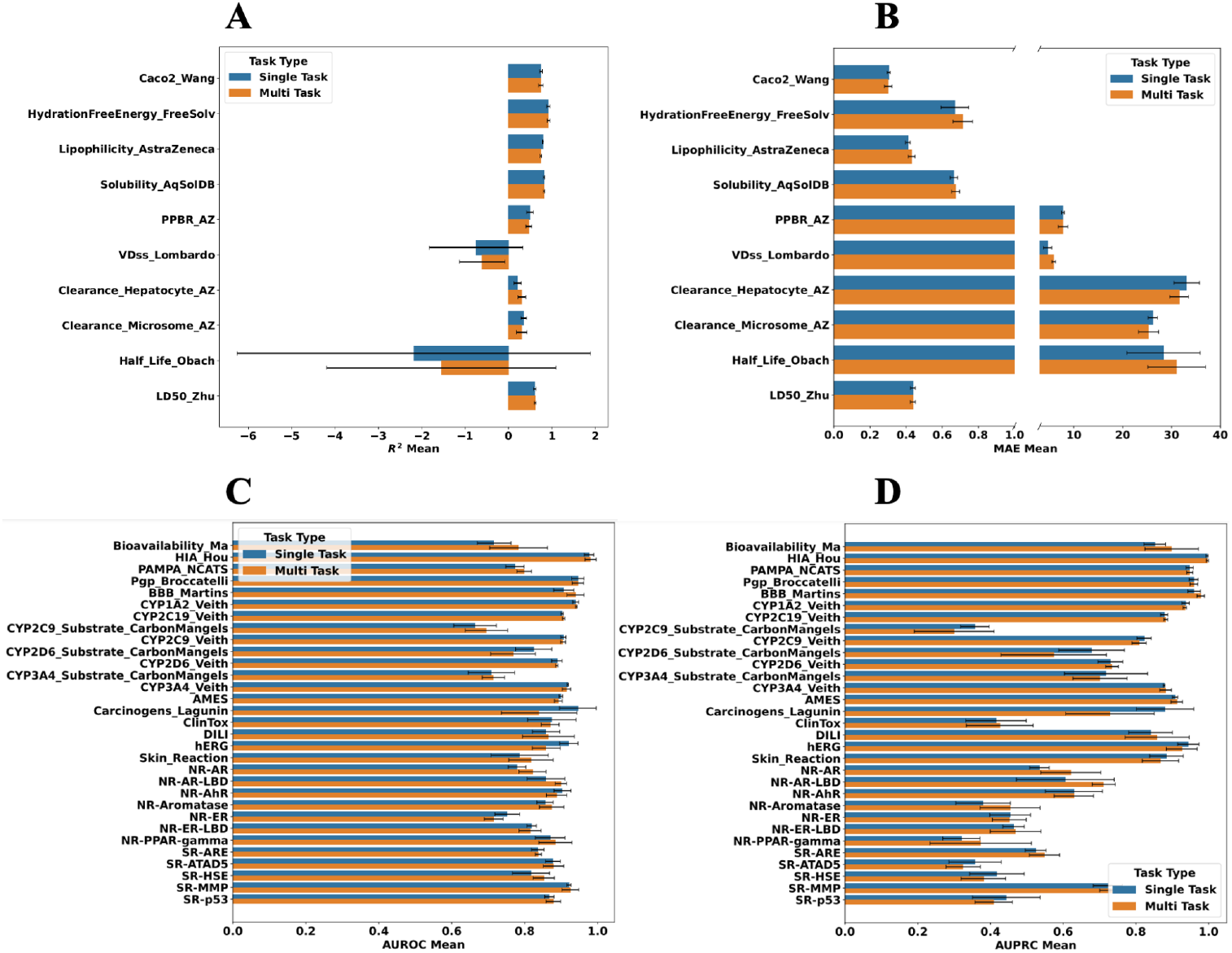
TDC Single-Task vs TDC Multi-Task Results. Performance of ADMET-AI on the 41 TDC ADMET datasets, trained either as 41 single-task models or as two multi-task models, one for regression and one for classification. **A)** Performance on regression datasets using the coefficient of determination (R^2^). **B)** Performance on regression datasets using mean absolute error (MAE). **C)** Performance on classification datasets using area under the receiver operating characteristic curve (AUROC). **D)** Performance on classification datasets using area under the precision-recall curve (AUPRC).

**Supplementary Figure 3:**
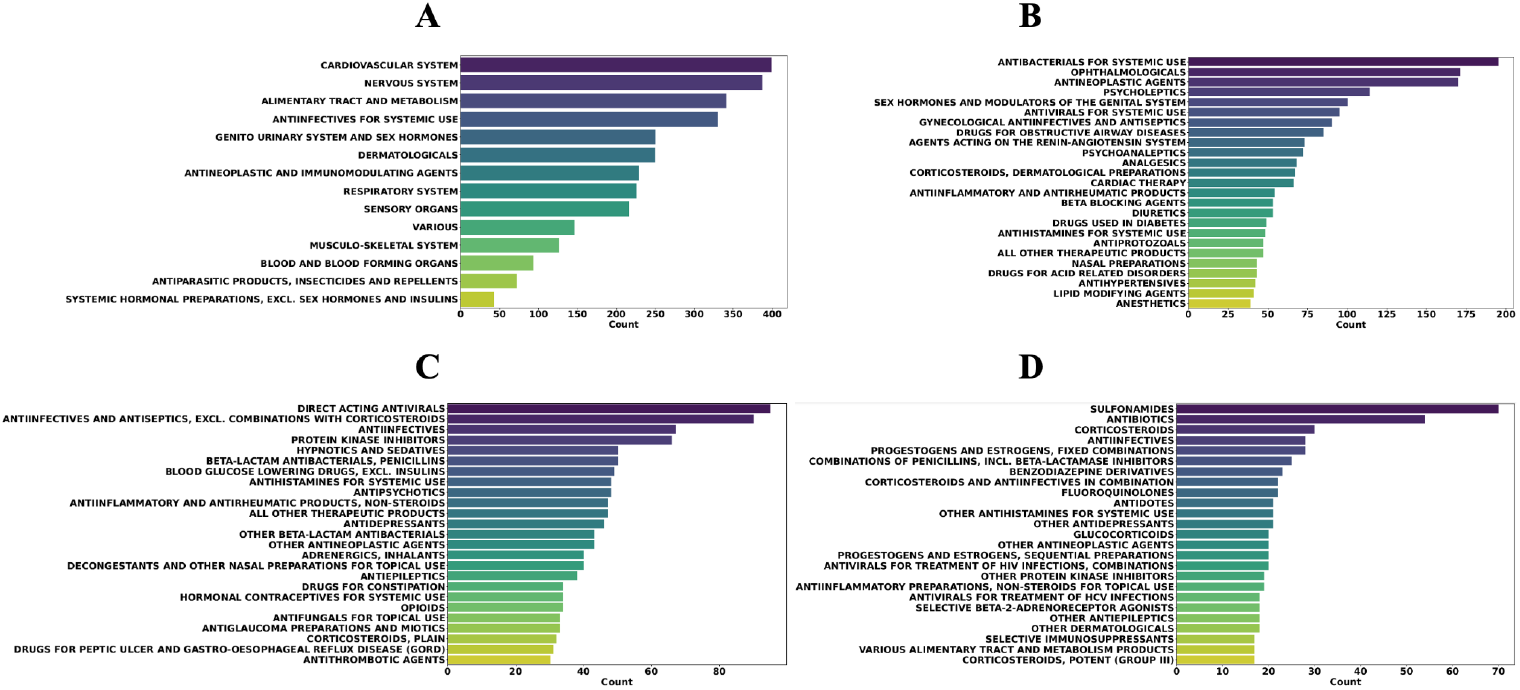
DrugBank ATC Codes. Frequencies of anatomical therapeutic chemical (ATC) codes in the 2,579 drugs in the DrugBank reference set at different ATC levels. **A)** The frequencies of the 14 ATC codes at level 1. **B)** The frequencies of the most common 25 of the 86 ATC codes at level 2. **C)** The frequencies of the most common 25 of the 223 ATC codes at level 3. **D)** The frequencies of the most common 25 of the 625 ATC codes at level 4.

**Supplementary Table 1: TDC Data Summary Statistics and Results**.

[TDC ADMET Results](https://docs.google.com/spreadsheets/d/1bh9FEHqhbfHKF-Nxjad0Cpy2p5ztH__p0pijB43yc94/edit?usp=sharing)

**Supplementary Table 2: DrugBank Reference Set of Approved Molecules with ATC Codes**.

[DrugBank Approved](https://docs.google.com/spreadsheets/d/1uKjqHMrAjnB8X-53cXgclb5jwXTortFeOYvEkCmNsSY/edit?usp=sharing)

**Supplementary Table 3: ADMET Tools Speed Comparison**.

[ADMET Speed Comparison](https://docs.google.com/spreadsheets/d/1_bvDO73MvYb7wl7_kH6Vagonyi-EJ6H-u8KpwQQ9Srg/edit?usp=sharing)

## References

1. Lyu, J., Irwin, J. J. & Shoichet, B. K. Modeling the expansion of virtual screening libraries. Nat. Chem. Biol. 19, 712–718 (2023).

2. Jacobs, S. A. et al. Enabling rapid COVID-19 small molecule drug design through scalable deep learning of generative models. Int. J. High Perform. Comput. Appl. 35, 469–482 (2021).

3. Bender, B. J. et al. A practical guide to large-scale docking. Nat. Protoc. 16, 4799–4832 (2021).

4. Feinberg, E. N., Joshi, E., Pande, V. S. & Cheng, A. C. Improvement in ADMET Prediction with Multitask Deep Featurization. J. Med. Chem. 63, 8835–8848 (2020).

5. Huang, K. et al. Therapeutics Data Commons: Machine Learning Datasets and Tasks for Drug Discovery and Development. in Proceedings of the Neural Information Processing Systems Track on Datasets and Benchmarks (eds. Vanschoren, J. & Yeung, S.) vol. 1 (Curran, 2021).

6. Yang, K. et al. Analyzing Learned Molecular Representations for Property Prediction. J. Chem. Inf. Model. 59, 3370–3388 (2019).

7. RDKit: Open-source cheminformatics.

8. Ali, K. M. & Pazzani, M. J. Error reduction through learning multiple descriptions. Mach. Learn. 24, 173–202 (1996).

9. Flask.

10. Wishart, D. S. et al. DrugBank 5.0: a major update to the DrugBank database for 2018. Nucleic Acids Res. 46, D1074–D1082 (2018).

11. WHO Collaborating Centre for Drug Statistics Methodology. https://www.whocc.no/atc/structure_and_principles/. ATC: Structure and Principles (2022).

12. Daina, A., Michielin, O. & Zoete, V. SwissADME: a free web tool to evaluate pharmacokinetics, drug-likeness and medicinal chemistry friendliness of small molecules. Sci. Rep. 7, 42717 (2017).

13. Xiong, G. et al. ADMETlab 2.0: an integrated online platform for accurate and comprehensive predictions of ADMET properties. Nucleic Acids Res. 49, W5–W14 (2021).

14. Pires, D. E. V., Blundell, T. L. & Ascher, D. B. pkCSM: Predicting Small-Molecule Pharmacokinetic and Toxicity Properties Using Graph-Based Signatures. J. Med. Chem. 58, 4066–4072 (2015).

15. Schyman, P., Liu, R., Desai, V. & Wallqvist, A. vNN Web Server for ADMET Predictions. Front. Pharmacol. 8, 889 (2017).

16. Tian, H., Ketkar, R. & Tao, P. ADMETboost: a web server for accurate ADMET prediction. J. Mol. Model. 28, 408 (2022).

17. Lee, S. K., Lee, I. H., Chang, H. J., Chung, J. E. & No, K. T. The PreADME Approach: Web-based program for rapid prediction of physico-chemical, drug absorption and drug-like properties. EuroQSAR 2002 Des. Drugs Crop Prot. Process. Probl. Solut. 418–420 (2003).

18. Lee, S. K. et al. The PreADME: PC-based program for batch prediction of ADME properties. EuroQSAR 2004 Des. Drugs Crop Prot. Process. Probl. Solut. 9–10 (2004).

